# Combination of oncolytic Maraba virus with immune checkpoint blockade overcomes therapy resistance in an immunologically cold model of advanced melanoma with dysfunctional T cell receptor signalling

**DOI:** 10.1101/2024.04.02.587705

**Authors:** Edward Armstrong, Matthew K.L. Chiu, Shane Foo, Lizzie Appleton, Pablo Nenclares, Anton Patrikeev, Nitya Mohan, Martin Mclaughlin, Galabina Bozhanova, Julia Hoebart, Victoria Roulstone, Emmanuel C Patin, Malin Pedersen, Joan Kyula, Fiona Errington-Mais, John C. Bell, Kevin J. Harrington, Alan A. Melcher, Victoria A. Jennings

## Abstract

**Background:** Over the past decade, cancer immunotherapies have revolutionised the treatment of melanoma; however, responses vary across patient populations. Recently, baseline tumour size has been identified as an independent prognostic factor for overall survival in melanoma patients receiving immune checkpoint inhibitors (ICIs). MG1 is a novel oncolytic agent with broad tumour tropism that has recently entered early phase clinical trials. The aim of this study was to characterise T cell responses in human and mouse melanoma models following MG1 treatment and to establish if features of the tumour immune microenvironment (TIME) at two distinct tumour burdens would impact the efficacy of oncolytic virotherapy.

**Methods:** Human 3D *in vitro* priming assays were performed to measure anti-tumour and anti-viral T cell responses following MG1 infection. TCR sequencing, T2 killing assay, and peptide recall assays were used to assess the evolution of the TCR repertoire, and measure specific T cell responses, respectively. *In vivo*, subcutaneous 4434 melanomas were characterised using RNAseq, immunohistochemistry (IHC), and flow cytometry. The effectiveness of intra-tumoural MG1 was assessed in advancing 4434 tumours and the generation of anti-tumour and anti-viral T cells measured by splenocyte recall assays. Finally, combination MG1 and α-PD-1 therapy was investigated in advanced 4434 tumours.

**Results:** MG1 effectively primed functional cytotoxic T cells (CTLs) against tumour associated antigens (TAA) as well as virus-derived peptides, as assessed using peptide recall and T2 killing assays, respectively. TCR sequencing revealed that MG1-primed CTL comprised larger clusters of similar CDR3 amino acid sequences compared to controls. *In vivo* testing of MG1 demonstrated that MG1 monotherapy was highly effective at treating early disease, resulting in 90% cures; however, the efficacy of MG1 reduced as the disease burden (local tumour size) increased, and the addition of α-PD-1 was required to overcome resistance in more advanced disease. Differential gene expression profiles revealed that increased tumour burden was associated with an immunologically colder TIME. Furthermore, analysis of TCR signalling in advancing tumours demonstrated a different dynamic of TCR engagement compared to smaller tumours, in particular a shift in antigen recognition by CD4+ cells, from conventional to regulatory subset.

**Conclusion:** Combination of MG1 with αPD-1 overcomes therapy resistance in an immunologically ‘cold’ model of advanced melanoma.

## Introduction

Oncolytic viruses (OVs) preferentially replicate within cancer cells, causing direct cytotoxicity, and induce both innate and adaptive anti-tumour immune responses. To date, a multitude of OVs have been tested in clinical trials and have been safe and well tolerated. The most clinically advanced agent, and the only virus currently approved for clinical use across the US, Europe, and Australasia, for local treatment of unresectable metastatic melanoma, is a genetically modified double- stranded DNA herpes simplex virus (HSV; JS-1 strain), talimogene laherparepvec (T-Vec; IMLYGIC®, Amgen Inc). The generation of local and systemic immune responses following oncolytic virotherapy has supported the rationale to combine OVs with other cancer immunotherapies, such as immune checkpoint inhibitors (ICIs). The MASTERKEY-265 (ClinicalTrials.gov: NCT02263508) trial, evaluated T-VEC in combination with pembrolizumab (αPD-1) for patients with advanced melanoma (stage IIIB-IVM1c). The Phase Ib part of this trial recruited 21 patients and confirmed that treatment was well tolerated, with no dose-limiting toxicities and encouraging early efficacy signals^1^. However, the full randomised, double blind, Phase III study (ClinicalTrials.gov: NCT02263508) was stopped early due to clinical futility, perhaps because outcome in the patient population tested was too good with pembrolizumab monotherapy for the addition of OV to make a significant difference, and/or because the protocol was altered between the early and later stages of the trial^2^.

Maraba virus was first developed as an oncolytic agent in 2010, and is a single-stranded, negative-sense, enveloped RNA virus that derives from the vesiculovirus genus of the Rhabdoviridae. Genetic modifications to the wild-type virus have resulted in the development of MG1, which has an enhanced capacity to replicate within tumour cells, a superior propensity to induce cancer cell death^3^ and has recently entered early clinical testing ^4,5^. MG1 has shown both oncotropic and cytotoxic activity in a range of murine and human cell lines. Pre-clinical studies have shown the successful application of MG1 as (i) a monotherapy^6^, (ii) a cancer vaccine vector expressing either tumour associated or viral antigens^7–9^, (iii) in combination with standard of care chemotherapeutic agents^10^ and (iv) in combination with ICI in a neoadjuvant setting^11^. All these features support the potential use of MG1 as an immunogenic oncolytic viral therapeutic agent. Despite a significant amount of pre-clinical data supporting MG1 as potent oncolytic agent capable of generating anti-tumour immunity in murine cancer models, to date there is limited data on the ability of MG1 to support the generation of human anti-tumour T cell responses, and what impact tumour size has on anti-tumour immune activation by MG1 or indeed other OVs. Therefore, the aim of this study was to monitor the generation and magnitude of anti-tumour and anti-viral T cell responses following MG1 treatment in both human and murine melanoma pre-clinical models. To do this we first used a human *in vitro* 3D melanoma T cell priming assays and measured primed cytotoxic T cells (CTL) responses against melanoma associated antigens, and against specific HLA-A2-restricted viral peptides. In addition, we performed T cell receptor (TCR) sequencing to track the evolution of the human TCR repertoire during MG1-induced CTL priming assays. In mice, using an early- and late-stage disease model of melanoma, we demonstrated that tumour size impacts on the TIME, consistent with other data with differing disease burdens^12,13^, and that a further feature of dysfunctional immunity in larger tumours is differences in CD4+ TCR engagement with antigen. MG1 monotherapy activated the TIME and was effective against small tumours generating long lasting anti-tumour immunity. For advance disease, however, the addition of αPD-1 therapy to MG1 was required to result in a significant survival benefit.

## Results

### MG1 infects, replicates in, and is cytotoxic against, melanoma cell lines grown either as 2D or 3D cultures

Previous studies have tested MG1 oncolytic activity against five human melanoma cell lines from the NCI60 (US National Cancer Institute) including M14, MALME3M, SKMEL28, UACC257 and UACC6 and three murine melanoma cell lines, B16, B16F10 and B16lacZ ^3,7,14^. To expand on this work, we investigated the oncolytic activity of MG1 in a further four human melanoma cell lines (A375, MEWO, MEL888 and MEL624) and the 4434 murine melanoma cell line. A dose and time dependent cytotoxicity was observed in all four human cell lines (Figure 1A) and in the murine 4434 cell line (Figure 1B) when grown as 2D monolayers. However, as 3D cultures better mimic the physical and biochemical features of a solid tumour mass, the ability of MG1 to infect, replicate in, and kill human melanoma cells grown in 3D cultures was next investigated. MEL888, MEWO and MEL624 cells all grew as 3D spheroid cultures and were successfully infected with MG1-GFP (Figure 1C). Viral replication (Figure 1D) and cytotoxicity (Figure 1E) were observed in all three cell lines tested. Hence, MG1 retains its oncolytic activity in tumour cells grown in 3D cultures.

**Figure 1:**
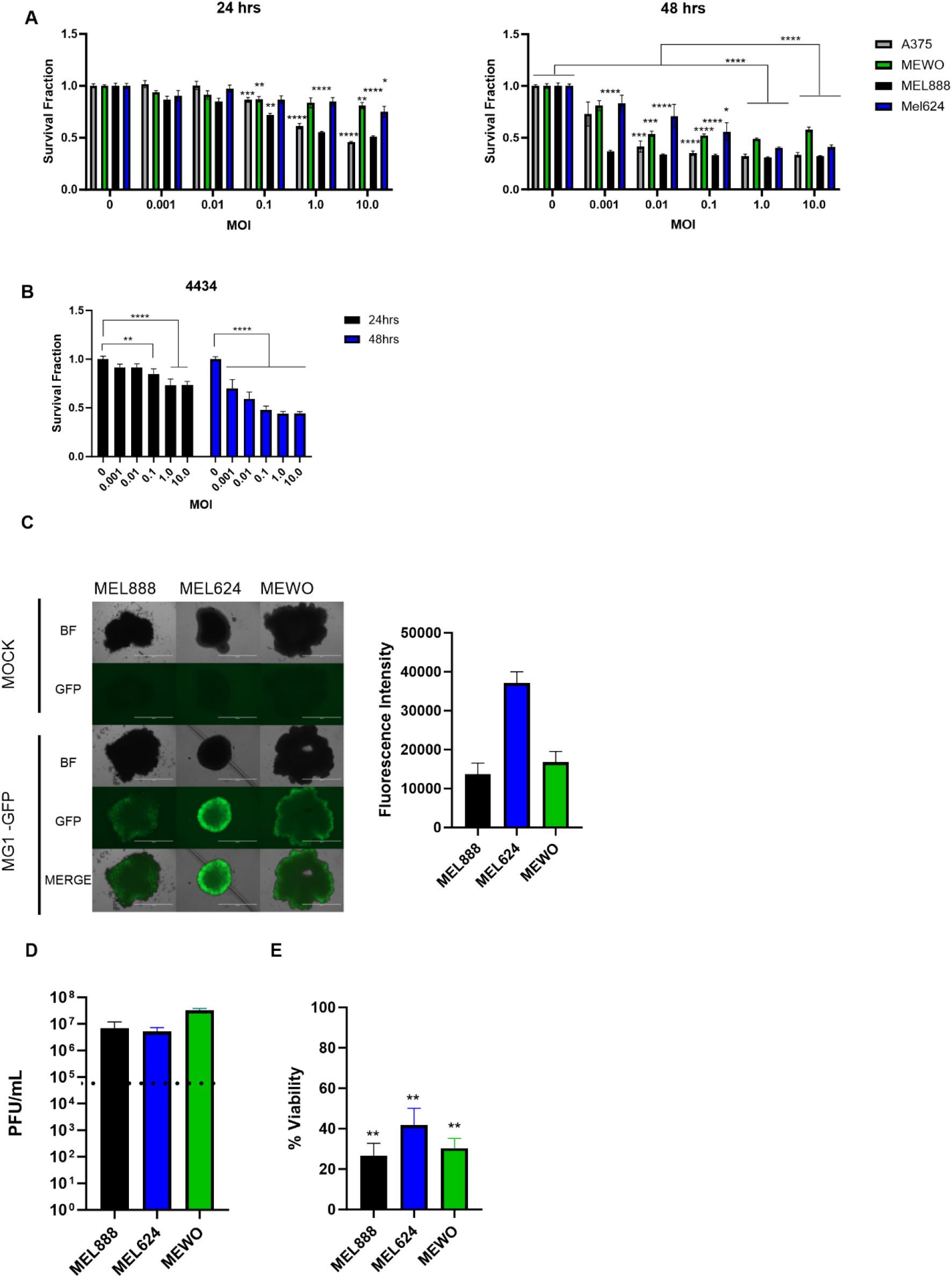
MG-1 infects, replicates and is cytotoxic against melanoma cell lines grown either in 2D or 3D cultures. Human **(A)** and murine **(B)** melanoma cell lines were treated with MG-GFP at concentrations ranging from 0 to 1pfu/cell, cell viability was determined by MTT assay at 24 and 48 hrs. Data shown is the average of three independent experiments ±SEM. **C/D and E.** Human melanoma tumour spheroids were infected with MG-GFP at an MOI 0.1, bright field and fluorescence images were taken and quantified at 20 hrs (C). Viral replication was determined at 24 hrs, input doses indicated with dotted line (D) and viability measured at 48 hrs (E). Data shown is the average of three independent experiments +SEM.

### MG1 primes specific anti-tumour and -viral T cell responses

We have previously developed *in vitro*, pre-clinical assays, to test the potential of OVs to support the activation of human adaptive anti-tumour immune priming using infected 2D monolayers as the melanoma ‘antigen source’, loaded on to immature dendritic cells (iDC) as antigen presenting cells (APC), for subsequent co-culture with responder T cells, to generate CTLs^15^. To increase clinical relevance, we adapted our model system to better mimic human disease and developed a 3D *in vitro* immune priming assay to test OV-induced anti-tumour immune priming^16^. To use 3D tumours as the ‘antigen load’, MEL888 spheroids were first infected with MG1 and cultured with iDC for 24 hrs; cell-free supernatants were then collected and assessed for a range of anti-viral, pro-inflammatory and immunosuppressive cytokines. MG1 infection induced the production of IFN-α, IL-28, IL-29, TNF-α, IP-10, although the difference with and without virus infection only reached significance for IFN-α, while levels of the immunosuppressive cytokine IL-10 remained unaltered (Figure 2A). As DC maturation is critical for effective T cell priming, the ability of MG1 to directly induce iDC maturation was next investigated. iDCs cultured with MG1 significantly increased cell surface expression of co-stimulatory molecules CD80 and CD86 (Figure 2B). To assess whether this MG1 induced DC phenotype, and pro-inflammatory cytokine production, supported adaptive immune priming, we tested MG1 infected MEL888 spheroids in CTL priming assays and showed that virus infection increased the production of melanoma-specific tumour associated antigen (TAA) T cell responses (Figure 2C). In addition, since anti-TAA priming has not previously been compared to anti-OV human T cell priming, we also investigated the generation of anti-viral responses under the same CTL priming conditions. NetMHCPan 4.1 was used to predicted the most immunogenic HLA-A2-restricted peptides in the MG1 proteins: M, G, N and P. T2 binding assays were then performed to confirm the ability of these 8-9mers to bind HLA-A2; the M protein peptide, RLGPTPPML, and the G protein peptide, SLIQDVERI, demonstrated the greatest significance in their ability to stabilise HLA-A2 on the surface of T2 cells (Figure 2D), indicating strong binding of these two peptides to the HLA-A2 molecule. To test the generation of anti-viral CTLs targeting these HLA-A2 restricted virus peptides, MEL624 (HLA-A2+) spheroids, with or without MG1 infection, were loaded onto HLA-A2+ iDC, and CTL priming assays were performed, as for the generation of anti-TAA responses. Primed CTL demonstrated increased cytotoxicity against RLGPTPPML and SLIQDVERI peptide loaded T2 cells compared to unloaded controls, reaching significance for RLGPTPPML (Figure 2E). Peptide recall responses against TYR overlapping peptide pools were also performed to confirm effective anti-tumour immune priming alongside anti-viral T cell responses (Figure 2F). Finally, TCR sequencing was performed on these primed CTL to explore the evolution of the TCR repertoire during T cell priming. TCR sequences were tracked between week one (one stimulation) to week two (two stimulations) from two donors (D1 and D2), to determine whether individual T cell clones were expanding or contracting. TCRs were clustered based on CDR3 amino acid triplet similarity using a kernel matrix. Week-2 clonotypes were then classified according to their normalised expansion rate ((frequency at week- 2/singleton frequency at week-2)/ (frequency at week-1)/singleton frequency at week-1) as expanded (if normalised expansion rate > 1) or contracted (if normalised expansion rate <1). Expansion and contraction of TCR clonotypes could be detected in both MG1- and mock- primed CTL. Interestingly however, clustering analysis of expanded clonotypes revealed that MG1-primed CTL, comprised of larger clusters of similar TCR sequences compared to the mock samples (Figure 2G, average cluster size in expanded clonotypes= 21.5 vs. 4.1), potentially representing the development of larger clusters of related T cell clonotypes targeting the virus and/or TAA specific antigens. Taken together, these data show for the first time that MG1 can support priming of human anti-tumour T cell responses, that the TCR repertoire of CTL during priming can be tracked and characterised, and that the functionality of the primed CTL can be measured using virus as well as TAA peptide recall assays.

**Figure 2:**
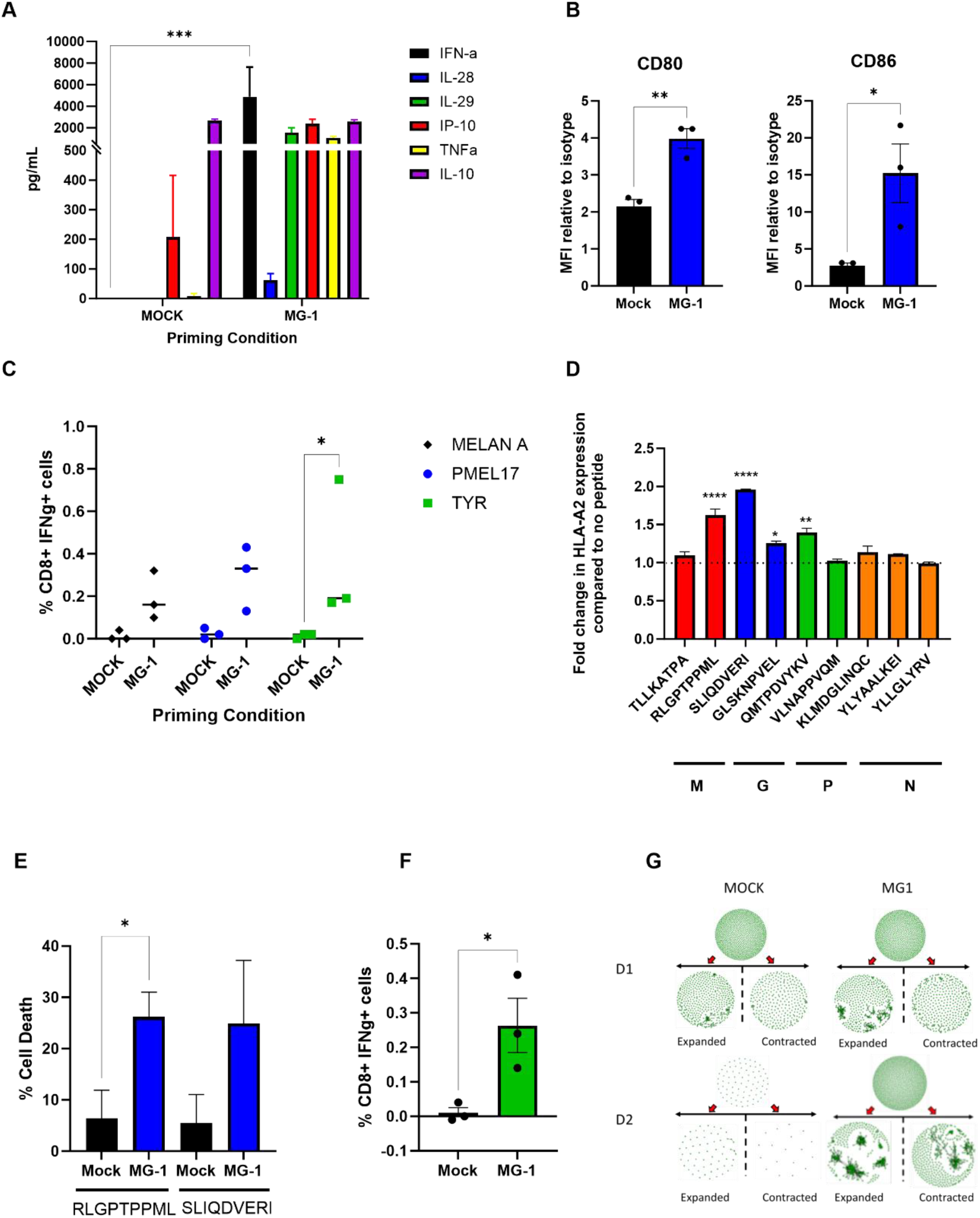
MG1 primes specific anti-tumour T cell responses. **A.** Supernatants from MEL888 spheroids cells treated with ±MG-1 and cultured with immature dendritic cells (iDC) for 24 hrs were collected and concentrations of IFNa, IL-28, IL-29, IP-10, TNFa and IL-10 were determined by ELISA. Data shows the mean +SEM from three independent experiments. **B.** iDC were treated with ±MG1 for 48h, cell surface expression of CD80 and CD86 was determined by flow cytometry. The mean fold increase in expression compared to isotype controls +SEM are shown (n=3). **C.** MEL888 spheroids were treated ±MG1 and cultured with iDC for 24 hrs before single cell suspensions were removed and cultured with autologous PBMC. CTL were re- stimulated once more then used in peptide recall assays against melanoma TAAs. The mean percentage CD8 cells expressing IFNg from three donors are shown. **D.** T2 cells were incubated with indicated MG1 peptides then cell surface expression of HLA-A2 was determined by flow cytometry, mean fold change in expression compared to no peptide control +SEM are shown (n=2). **E and F** MEL624 spheroids were treated ±MG1 and CTL generated as described in C **E.** 4hr Cr^51^ killing assay against T2 cells loaded with indicated peptides. **F.** Peptide recall assay against TYR, mean percentage of CD8 cell expressing IFNγ from three donors are shown. **G** Network diagrams of CDR3β amino acid clustering at week-1 (top) and week-2 expanded and contracted repertoires based on normalised expansion rate for each donor (D1 and D2). Each circle within the network represents a TCR clone and each cluster is represented by a group of TCRs linked by vectors.

### Advanced 4434 tumours are immunologically colder than early disease

Whilst it is well-recognised that immunotherapy is more effective in patients with a lower tumour burden^17,18^, the evolution of the immune microenvironment as tumours grow, and how this impacts the efficacy of immune-based treatments, is poorly understood. Therefore, prior to testing MG1 *in vivo*, we first investigated the transcriptional and phenotypical differences between small (50mm^3^) and large (150mm^3^) 4434 melanomas growing in C57BL/6 mice. Bulk RNA sequencing was performed on small and large tumours (n=3 mice per group). Differential gene expression revealed that large tumours had 589 genes upregulated and 1015 genes downregulated (by greater than 2-fold, p<0.05) compared to smaller tumours (Figure 3A). To investigate which biological processes were changing in large tumours gene ontology (GO) analysis was performed on both up- and down-regulated genes. GO processes that were downregulated in large tumours highlighted multiple immunological pathways, such as inflammatory response, T cell activation and positive regulation of cytokine production (Figure 3B). As multiple immune pathways were identified to be dysregulated in larger tumours, we next assessed the total immune and stromal composition of both small and large 4434 tumours using the mMCP counter tool; total T, CD8+, NK, B and myeloid cells were reduced in large tumours compared to small (Figure 3C). To validate these changes in the immune composition, the number of CD8+ cells was assessed by IHC and flow cytometry analysis. Immunohistochemical and FACS-based staining confirmed that within large tumours, CD8+ cells were reduced, both as an individual population (per mm^2^) and as a proportion of total T cells (Figure 3D and E, respectively). Moreover, the number of FOXP3+ CD4+ cells increased as a proportion of the total CD4+ T cell population in larger tumours (Figure 3E). To further interrogate the differences in the transcriptomics of small and large tumours, we focused on the expression of a subset of immune-specific genes (Figure 3F). Small tumours displayed an increased expression of genes involved in antigen presentation (*H2-K1*, *H2-D1*, *H2-M3*, *H2-T23*, *H2-Aa*, *H2-Eb1*, *Cd74*), co-stimulation (*Icos*, *Cd27*, *Cd28*, *Tnfrsf4*, *Tnfsf14*, *Cd40*) and immune cell recruitment (*Ccl2*, *Ccl5*, *Ccl7*, *Cxcl9* and *Cxcl10*) compared to large tumours. Taken together, small 4434 tumours are immunogenically ‘hotter’ than more advanced larger tumours, which display a reduced immune cell infiltrate and reduced expression in key genes involved in anti-tumour immune responses.

**Figure 3:**
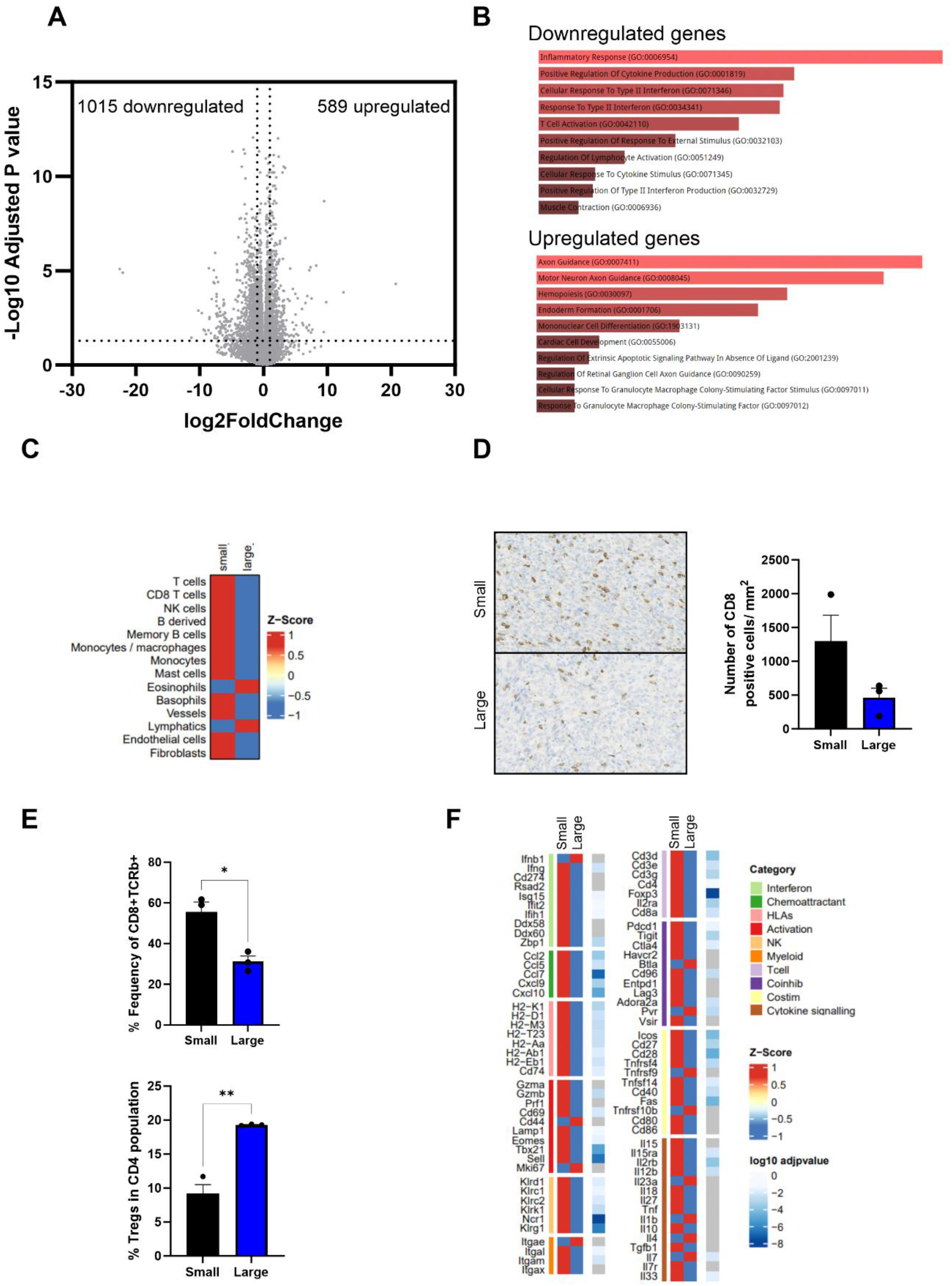
Advanced 4434 tumours are immunologically colder than early disease. 4434 cells (4x10^6^) were injected s.c. into C57BL6 mice, tumours were collected when they reached an average volume of either 50mm^3^ (small) or 150mm^3^ (large) (n=3 per group). **A.** Shows the volcano plot of differentially expressed genes in small vs large tumours. The x-axis and y-axis are the log2(fold change) and -log10(p- adjusted) values, respectively (dotted lines indicate 2-fold change on x-axis and p<0.05 on y-axis). **B.** Shows the top-10 differentially enriched GO biological processes (adjusted p<0.05) that are associated with both up and downregulated genes sets. **C.** Heatmap showing Z-score normalised mMCP counter scores in small and large tumours. **D.** Immunohistochemical analysis of CD8 expression in small- (top) and large-tumours (bottom), quantification of number of positive CD8/mm^2^ was determined by QuPath, mean +SEM is plotted. **E.** Tumours were homogenised and cell surface expression and intracellular staining of TCRb, CD8, CD4, CD25 and FoxP3 were analysed by flow cytometry. **F.** Heatmaps showing Z-scores of normalised immune gene expression in small and large tumours.

### T cell receptor dynamics differs in more advanced tumours

A further characteristic of tumours is the dynamics of their TCR signalling, which we have previously shown to impact on oncolytic virus therapy^19^. Therefore, to further evaluate differences in T cell function between small and large tumours, tumours from 4434-bearing Nr4a3-Tocky mice that exhibited either limited (<50mm^3^, small) or enhanced growth (>100mm^3^, large) at 21 days post-implantation, were analysed by FACs. The Tocky model is a transgenic mouse *in vivo* system that incorporates an unstable fluorescent reporter protein in the promoter region for Nr4a3, an intermediate-early gene downstream of TCR signalling^20^. Upon TCR signalling, Nr4a3 is transcribed resulting in blue fluorescence, which decays over time to red with a half-life of 6 hours. If the TCR is persistently engaged, and Nr4a3 continually transcribed, new blue fluorescence is seen within the cell in addition to decaying red, resulting in blue/red positive ‘persistent’ T-cells (Figure 4A shows an illustrative flow cytometry schematic). This model enables isolation and analysis of the antigen-reactive T-cell population within a tumour *in vivo* with a temporal component, and in this context provides valuable information regarding the occupancy of the ‘antigen niche’ within the TME as growing tumours escape immune control.

**Figure 4:**
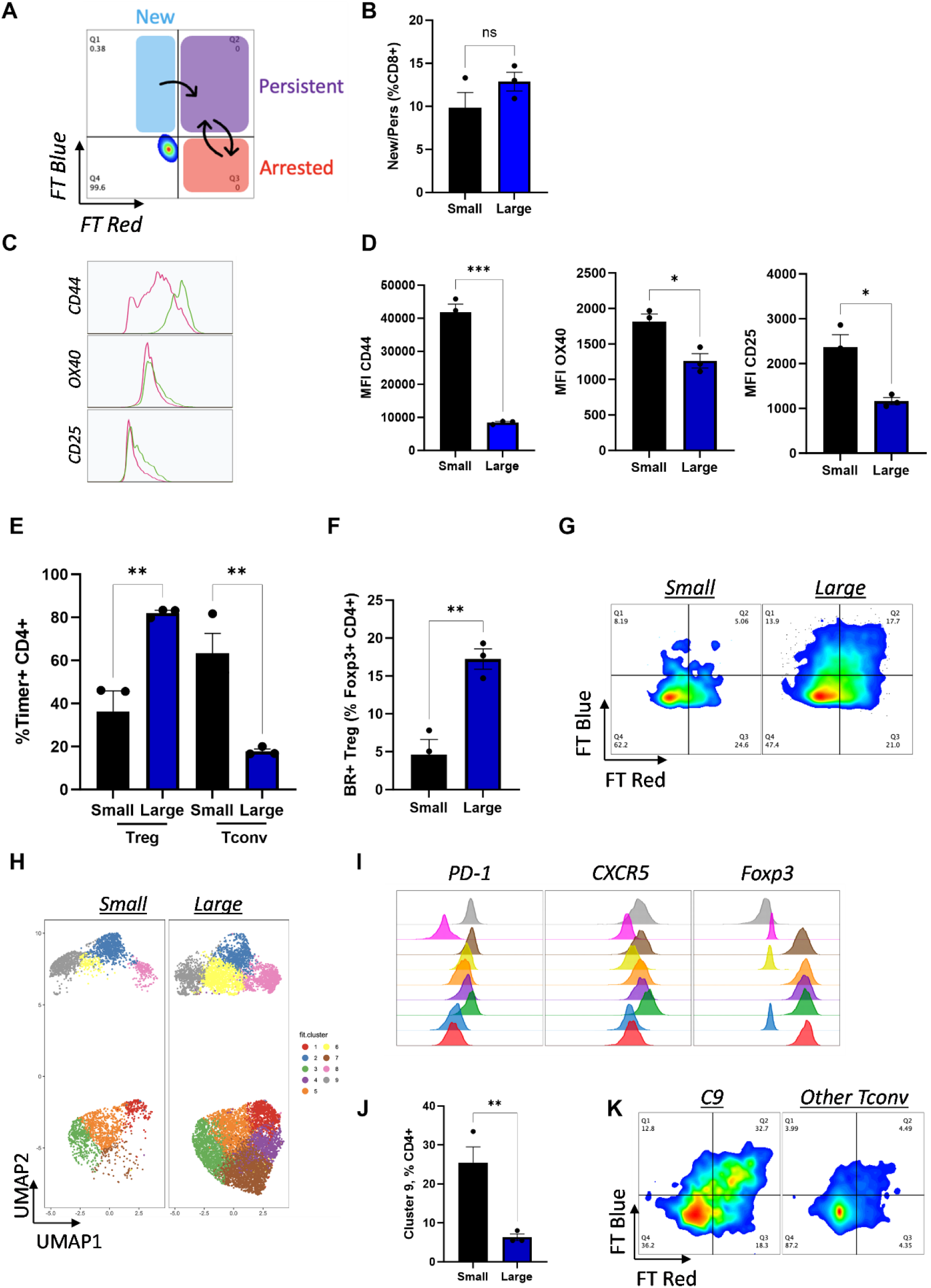
Analysis of the dynamics of antigen engagement in large and small tumours. 4434 cells (4x10^6^) were injected s.c. into Nr4a3 Tocky mice, tumours were collected when they reached an average volume of either 50mm^3^ or >100mm^3^ (n=3 per group) homogenised and analysed by flow cytometry. **A** Schematic of Tocky fluorescence following antigen engagement, “New” blue+ (B+), “Persistent” Blue+/Red+ (BR+) and “Arrested” Red+ (R+). **B** The percentage of recently engaged (B+/BR+) CD8+ T cells. Expression of CD44, OX40 and CD25 on CD8+ T cells with recent antigen engagement. **C** Representative histograms (Small=green and Large=pink) and **D** mean fluorescence intensity values. **E** The percentage of Timer+ CD4+ cells in Tconv and Treg T cell subsets. The percentage of BR+ Treg in total Treg CD4+FoxP3+ populations **F** representative flow plot shown in **G. H)** UMAP cluster analysis of the CD4+ TIL population. **I** Percentage of cluster 9 as a percent of CD4+ cells. **J** Representative histograms showing expression of PD-1, CXCR5 and FoxP3 on CD4+ TIL clusters. **K** Tocky fluorescence of cluster 9 (C9) compared to other Tconv clusters in small tumours.

When comparing CD8+ T cells within small and large tumours, although no change in the absolute frequency of recently antigen-engaged cells (Tocky Timer Blue+/BlueRed+) was demonstrated between small and large tumours (Figure 4B), TCR-engaged CD8+ T cells within small tumours had significantly higher expression of the T-cell activation/memory markers CD44 and CD25, and the co-stimulatory receptor OX40 (Figure 4C, D), suggesting a better CD8+ T cell fitness response to antigen-engagement.

Within the CD4+ compartment, a shift was seen in the composition of the total antigen-reactive population (Tocky Timer + CD4+) between small and large tumours. In large tumours, this population was primarily composed of regulatory T-cells (CD4+ FoxP3+ Treg) (Figure 4E), which had high levels of persistent antigen-engagement (Figure 4F, G), a characteristic of effector Treg ^21^. In contrast, CD4+ FoxP3- conventional CD4 (Tconv) predominated within the antigen-reactive CD4+ population in small tumours (Figure 4E), suggesting a more supportive, less immunosuppressive environment for therapy in smaller tumours. UMAP cluster analysis of tumour-infiltrating CD4+ T cells further illustrates this shift (Figure 4H) and demonstrates a population of CD4+ Foxp3- Tconv that are significantly enriched in small tumours (cluster 9, Figure 4H, J). Marker analysis of this cluster reveals high expression of PD-1 and CXCR5 (Figure 4I), characteristic of follicular helper Tcells (Tfh). This cluster is highly antigen-engaged when compared to the other T-conv clusters (Figure 4K), suggesting Tfh may be instrumental in maintaining an immune response within small tumours. This subset has also been demonstrated to aid response to therapies targeting the PD-1/PD-L1 axis, although confirmation of their role in cancer remains to be elucidated^22,23^ .

#### A single intra-tumoural injection of MG-1 is highly successful in curing early disease burden but is ineffective in advanced disease

With the differences in TIME observed in early and late 4434 disease states, we decided to test how effective MG1 was in tumours that were small (50mm^3^), medium (100mm^3^), or large (150mm^3^). MG1 was highly effective at treating small tumours, curing 90% of animals (Figure 5A). Medium tumour-bearing mice were less responsive to MG1 treatment, with 40% of mice displaying long term survival; the median survival of this group was 77 days compared to 41 days in untreated mice (p < 0.0001) (Figure 5A). However, MG1-treated large tumour-bearing mice only showed a small increase in survival when compared to untreated mice (PBS median survival 41 days vs 46 days for large tumour MG1-treated tumours p=0.0486) (Figure 5A). To determine whether cured mice generated long-term anti-tumour immunity, successfully treated, cured mice were re-challenged with 4434 tumours alongside naïve control mice (Figure 5B). While naïve mice developed 4434 tumours, no tumour growth was observed in the cured mice, indicating that long term immunity had been generated following MG-1 treatment, regardless of the initial size of the tumour (small or medium). To understand the global immune effects of MG1 on small and large tumours, we used RNAseq to investigate the transcriptional changes occurring within the different sized tumour 48 hrs following viral injection. Following MG1 treatment, small tumours demonstrated 5037 significantly differentially expressed genes, while large tumours had only 485, indicating a greater impact on global gene expression in small tumours following MG1 (Figure 5C, D, respectively). The GO biological processes that were associated with upregulated genes in small tumours following MG1 treatment, included positive regulation of cytokine production, inflammatory response, and regulation of type II IFN and TNF. Conversely upregulated genes in large tumours treated with MG1 did not contain any of these processes; instead GO biological processes against virus infection predominated, including defence to virus, negative regulation to virus process and replication, and anti-viral innate immune response. Next, we investigated the response to MG1 treatment on our targeted subset of immune-specific genes. MG1- treated small tumours had increased expression of many genes associated with immune activation compared to untreated small, and MG1-treated large tumours, including antigen presentation, chemoattraction, IFN response and cytokine signalling, as well as expression patterns associated with greater immune cell infiltration and upregulation of costimulatory and coinhibitory molecules (Figure 5E). CD8 expression in small and large tumours following MG1 treatment was validated at the protein level by immunohistochemical staining; MG1- small- treated tumours demonstrated a significantly higher number of CD8 cells/mm^2^ when compared to MG1-large- treated tumours (Figure 5F). Finally, to measure the level of anti-tumour immunity generated following MG1 treatment, splenocytes were collected from medium- or large- MG1 treated tumour-bearing animals. Splenocytes from medium-MG1-treated animals displayed a recall response against 4434 tumour cells *ex vivo* as measured by IFNγ release, while splenocytes isolated from large-MG1-treated animals showed no such responses (Figure 5G). These results indicate that the magnitude of the immunological response to MG1, and the generation of tumour specific T cells is impeded in locally advanced, relative to earlier, disease, due to the more immunosuppressed TIME.

**Figure 5:**
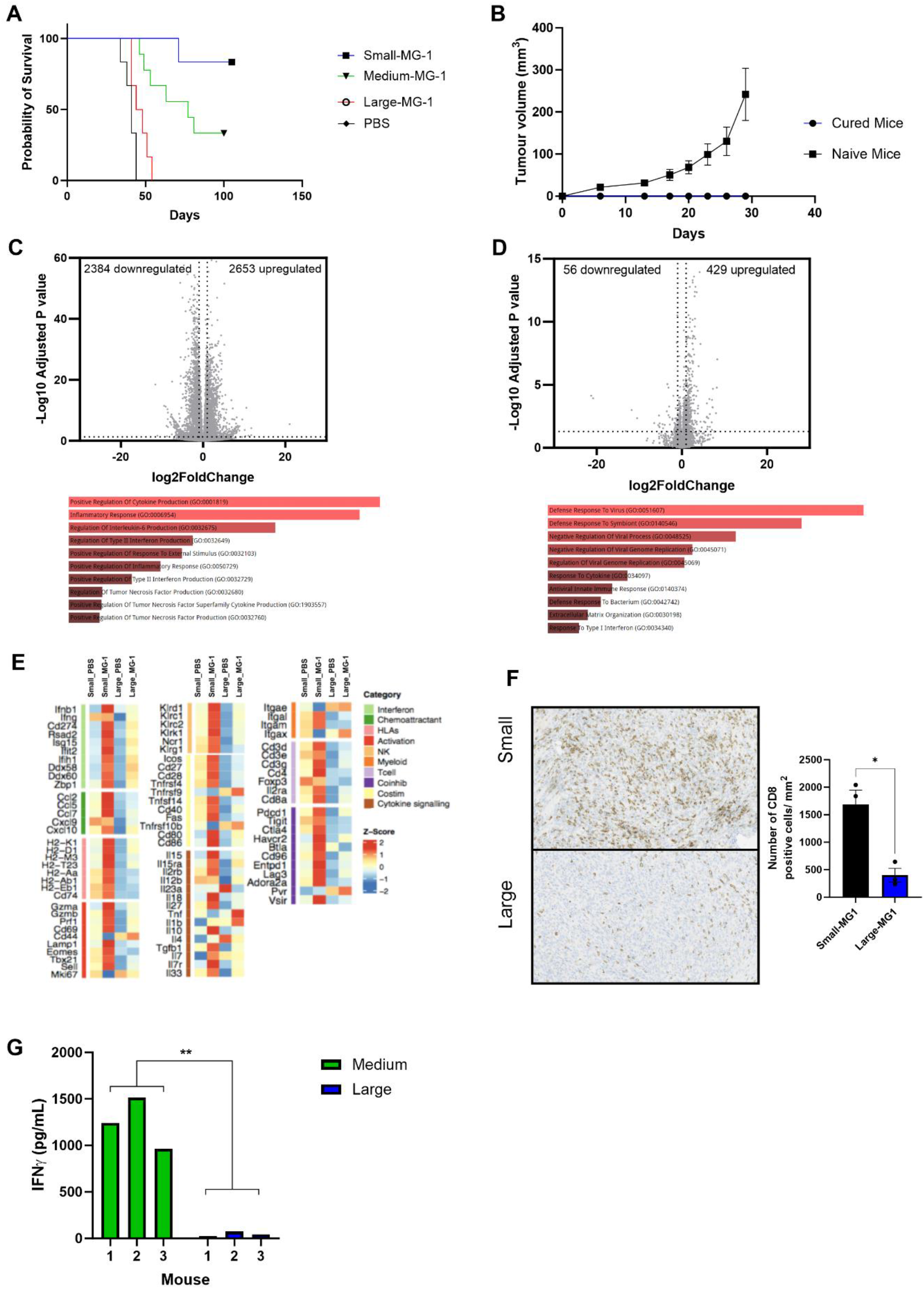
A single intra-tumoural injection of MG-1 is highly successful in curing early disease burden but is ineffective in advance disease. 4434 cells (4x10^6^) were injected s.c. into C57BL6 mice intra-tumoural injection of MG-1 (1x10^7^ PFU) or PBS were performed on small, medium, and large tumours (50-150mm^3^). **A.** Kaplien- Meier survival curve of tumour bearing mice (6 mice per group). **B.** Mice cured with MG1 treatment (5 mice) were injected with 4434 cells (4x10^6^) alongside naïve mice (6 mice) and growth of s.c. tumours plotted overtime. Graph shows the average tumour growth +/-SEM. **C and D**. Show the volcano plot of differently expressed genes following MG1 treatment of small (C) and large (D) tumours. The x-axis and y-axis are the log2 (fold change) and -log10(p-adjusted) values, respectively (dotted lines indicate 2-fold change on x-axis and p<0.05 on y-axis). Top-10 differentially enriched GO biological processes (adjusted p<0.05) that are associated with both up and downregulated genes are displayed underneath. **E.** Heatmaps showing Z-scores of normalised immune gene expression in small and large tumours ± MG1 treatment. **F.** Immunohistochemical analysis of CD8 expression in small (top) and large (bottom) 4434 tumours 48 hrs following MG1 treatment, quantification of number of positive CD8/mm^2^ was determined by QuPath, mean +SEM is plotted. **G.** Splenocytes from individual C57BL/6 mice bearing medium or large s.c.4434 tumours treated with MG1, were re-stimulated *in vitro* with 4434 tumour cells. Forty-eight hours later, supernatants were assayed for secretion of IFN-γ by enzyme-linked immunosorbent assay. Graphs show the concentration of IFN-γ from individual mice (3 mice/group).

### Combination of MG-1 with a-PD-1 improves anti-tumour immunity and survival compared to monotherapy in locally advanced disease

As MG1 as a monotherapy against more advanced disease did not generate long term cures, we hypothesised that the addition of αPD-1 to the treatment regimen would improve the generation and function of anti-tumour T cells. Importantly, PD-L1 (*Cd274*) and PD-1 (*Pdcd1*) were both upregulated following MG1 treatment (Figure 5E). This was confirmed at the protein level by immunohistochemical and FACS-based methods, which demonstrated a significantly increased expression of PD-L1 following MG1 treatment of both small and large tumours (Figure 6A and B).

**Figure 6:**
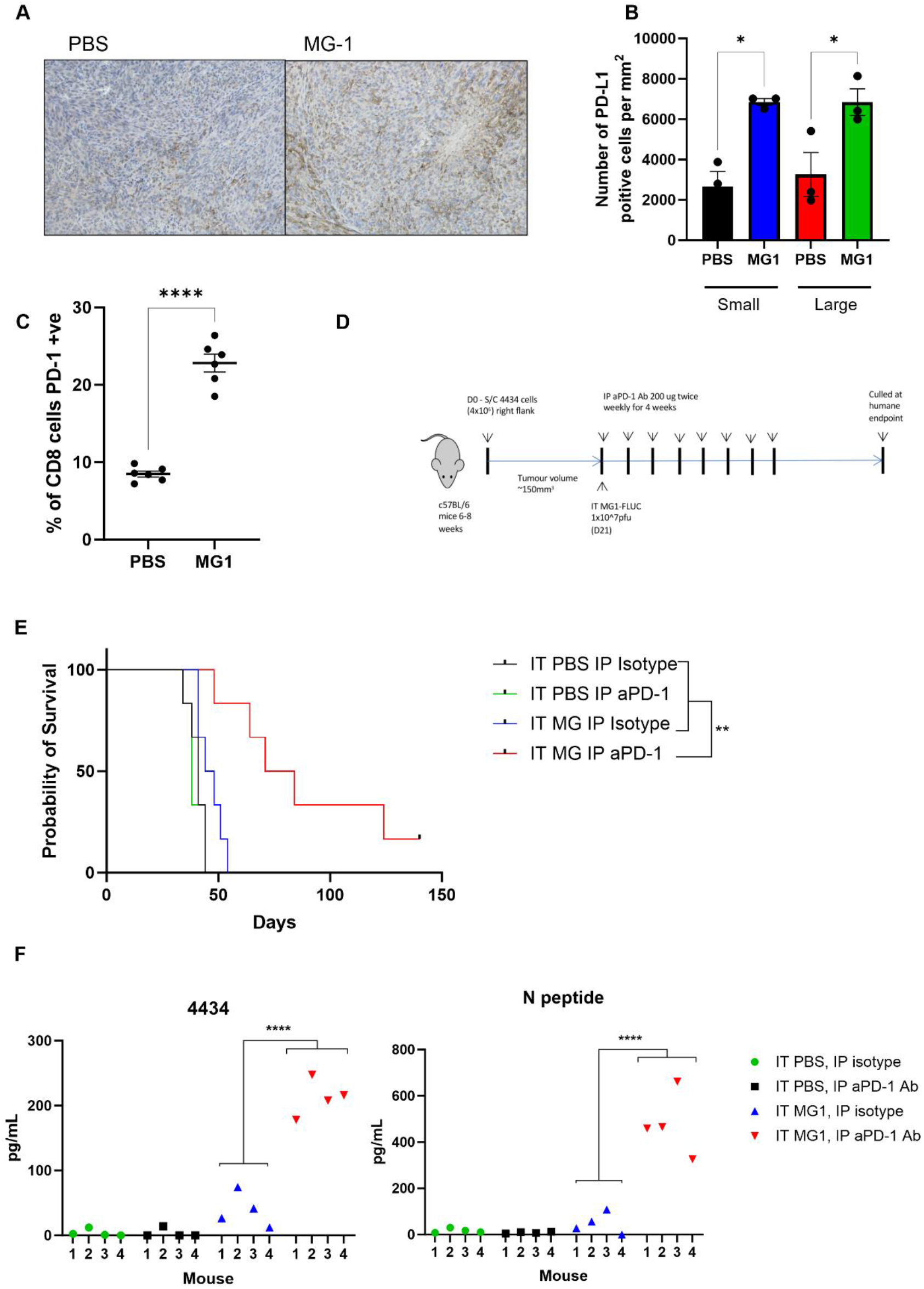
Combination therapy of MG-1 with anti-PD-1 overcomes resistance of monotherapies in larger tumours. **A.** PD-L1 expression was determined by immunohistochemistry in small and large tumours following MG-1 treatment (left, large-PBS treated; right large-MG1 treated). **B.** The number of positive PD-L1/mm^2^ was determined by QuPath, mean +/-SEM is plotted. **C.** Splenocytes were isolated from PBS (control) or MG-FLUC treated animals and PD-1 expression on CD8 T cells determined by flow cytometry. The mean percentage PD-1 expression on CD8 positive splenocytes is shown +/- SEM (5 mice per group). **D.** Schematic of treatment regime. **E** Kaplien-Meier survival curve of treated animals (6 mice/group). **F** Splenocytes from individual C57BL/6 mice bearing s.c.4434 tumours and treated with a combination of i.t. PBS or MG-1 and i.p. isotype control antibody or anti-PD-1 antibody as labelled, were re-stimulated *in vitro* with 4434 tumour cells or N peptide. Forty-eight hours later, supernatants were assayed for secretion of IFN-γ by enzyme- linked immunosorbent assay. Graphs shown the concentration of IFN-γ from individual mice (4 mice/ group

Furthermore, PD-1 expression was significantly increased on splenic CD8 T cells seven days following i.t MG1 (Figure 6C), indicating a systemic effect of local MG1 administration and thus implicating this axis as a valid target for combination immunotherapy. Large 4434 tumours were treated with either single agent alone (in combination with PBS or Isotype control) or MG1 and αPD-1 co-treatment. ICI was given twice weekly following a single MG1 i.t. injection (Figure 6D). MG1 and αPD-1 were ineffective when delivered as monotherapies in this advanced disease setting (median survival PBS/Isotype; 41 days, PBS/aPD-1; 38 days, MG1/Isotype; 46 days); however, the addition of αPD-1 to MG1 significantly increased survival (median survival 77.5 days) (Figure 6E). *Ex vivo* splenocyte recall assays were performed to assess the level of anti-tumour immunity generated following combination therapy, the addition of αPD-1 to MG1 treatment significantly enhanced priming against 4434 tumour cells (Figure 6F). In this experiment we also tested priming against MG1, by pulsing splenocytes with a defined H-2b restricted rhabdovirus N protein peptide; consistent with the human data from Figure 2E, activation of an immune response against tumour was accompanied by an anti-viral response, with both responses enhanced *in vivo* by the addition of αPD-1 to virotherapy (Figure 6F). Taken together, these results suggest that MG1 in combination with αPD-1 may be an effective treatment option particularly for more advanced melanoma, with partial reversal of the immunosuppressive microenvironment in larger tumours being boosted by the addition of ICI to reveal a therapeutic effect.

## Discussion

OVs are a promising cancer immunotherapy agent due to their direct lytic effect followed by the generation of anti-tumour T cell responses. The induction of an effective anti-tumour T cell response is critical to generating long-term responses following oncolytic virotherapy. Therefore, gaining a greater understanding of conditions required for effective T cell clearance of tumours and the induction of antigen specific T cell responses is essential to identify cancer patients that have the best chance of responding to oncolytic virotherapy, and to facilitate the design of combination therapy strategies. The evolution of the TIME as a tumour grows, and how this impacts the efficacy of immune-based treatments, is poorly understood, and has not been extensively investigated in murine models. Therefore, in this study we characterised 4434 tumours at different levels of disease burden to try and understand the impact of tumour growth on immune features in the TME and investigated how these changes may impact successful virotherapy. RNA sequencing revealed a significant change in immune composition during tumour growth, which was confirmed at the protein level with immunohistochemical staining and flow cytometry. We found that, as these tumours grew, they became immunological colder with fewer immune cells and a reduction in immune stimulatory genes; whilst such data may be unsurprising, it provides clear leads on what may be important targets to pursue in the more advanced disease setting. This pre-clinical data supports the clinical findings that large tumours are more immunosuppressive compared to small tumours^13,17^, directly impacting the ability of the host immune system to effectively mount a natural or immunotherapy-induced immune response. Although our study focussed on the immunosuppressive features of the local TIME, this doesn’t reflect the full nature of the immunosuppressive impact of increased tumour burden, which will be systemic as well as local^12,24^. Performing similar characterisation in other models and correlating these findings with human datasets will be important to validate shared targets of most relevance for testing in mouse models. Interestingly, whilst both αPD-1 therapy and MG1 were ineffective as single agents in advanced disease, combination treatment led to a significant extension of survival that was associated with an increase in both anti-tumour and anti-viral T cells. It is noteworthy that both our human and mouse systems show priming against tumour is accompanied by priming against virus; however, the importance and the relative contribution of the T cell response against the virus relative to the tumour for effective oncolytic virotherapy therapy remains unknown. The findings of the current study support the potential importance of the anti-viral response in initiating and/or maintaining an anti-tumour effect and highlight the relevance of tracking both these targets of immune priming as translational readouts within oncolytic virus trials.

Overall, this study shows that combining MG1 with αPD-1 therapy has the potential to overcome therapy resistance in an immunological ‘colder’ advanced tumour TME. Its broader implications highlight both the need to understand the biology of more advanced relative to earlier stage cancer at baseline, so that appropriate treatment targets can be selectively identified and pursued using single agent or combination strategies, and the value of using orthogonal both human and murine pre-clinical systems to maximise the impact of laboratory studies on the design and interpretation of clinical studies. Here, using these approaches, we have rationalised a single agent OV approach for the treatment of early melanoma, with immune checkpoint combination suitable and required only for more advanced disease. Finally, these findings throw light on negative immunotherapy trials in which the wrong stage of disease may have been targeted^2^. For example, when αPD-1 was added to oncolytic virotherapy in melanoma, the trial was designed to exclude those patients with the most advanced disease, who had not benefitted from single agent virus in a previous trial^25^. Our data suggests that the most advanced patients are those for whom the combination would nevertheless have been of greatest benefit, which may explain why no significant difference was seen with αPD-1 +/- OV in the study as designed^26^. Hence understanding and incorporation of disease stage into immunotherapy trials in pre-clinical models may have significant implications for studies as designed and delivered in the clinical setting.

## Material and Methods

### Cell culture and reagents

A375, MeWo, T2 and Vero cell lines were purchased from ATCC and authenticated using STR profiling and comparison with the DSMZ database. Mel-624 and Mel-888 were obtained from the Cancer Research UK cell bank. The BRAF-mutant (BRAFV600E) mouse melanoma cell line 4434 was established from C57BL/6_BRAF +/LSL-BRAFV600E; Tyr:CreERT2+/o^27^. All cell lines were grown in glutamine-containing DMEM (Sigma-Aldrich Ltd), supplemented with 10% FCS (*v*/*v*) (Sigma-Aldrich Ltd), apart from T2 cells which were grown in glutamine containing RPMI (Sigma-Aldrich Ltd), supplemented with 10% FCS (*v*/*v*). All cell lines were routinely checked for mycoplasma and were free from contamination.

PBMC were isolated from healthy donor volunteers after written, informed consent was obtained in accordance with local institutional ethics and review approval. PBMC were isolated from whole blood by density gradient centrifugation on Lymphoprep® (StemCell Technology) and cultured at 2x10^6^ cells/mL in glutamine containing RPMI, supplemented with 10% FCS (*v*/*v*). CD14^+^ cells were isolated from PBMC using MACS isolation procedures, following the manufacturers’ protocols (Miltenyi Biotec). Immature DC (iDC) were generated by culturing CD14+ cells in glutamine-containing RPMI supplemented with 10% FCS (*v/v*), recombinant human IL-4 500 IU/mL and GM-CSF 800 IU/mL (both R&D systems) at a cell density of 1-2 x10^6^ cells/mL for 5 days. Cytotoxic T lymphocytes (CTL) were cultured at 4-6x10^6^ cells/mL in in glutamine-containing RPMI supplemented with 7.5% (*v/v*) human AB serum, 1 mM sodium pyruvate, 1 mM HEPES; 1% (*v/v*) non-essential amino acids, 20 µM 2β- mercaptoethanol (all Sigma-Aldrich Ltd) and recombinant human IL-7 (5 ng/mL) (R&D Systems).

### Viruses

MG-1 expressing green fluorescent protein (MG-GFP) and firefly luciferase (MG- FLUC) were provided by Ottawa Hospital Research Institute and virus amplified and titre determined by standard plaque assay on Vero cells^3^.

### MTT Cell Viability

Melanoma cell lines were treated with MG-GFP at indicated doses for 24 and 48 hrs. 20µL MTT (5mg/mL; Sigma-Aldrich Ltd) was added to cells 4 hrs prior to the end of the incubation period. After 4 hrs, tissue culture supernatant was removed, and cells were solubilised using 150µl DMSO (Sigma-Aldrich Ltd). Optical density absorbance readings were determined using a Thermo Multiskan EX plate reader (Thermo Fisher Scientific), at 540 nm absorbance.

### Infection of tumour spheroids

Mel-888, MeWo and Mel-624 cells were seeded into 96-well ultra-low binding plates (Corning®) at a density of 2.5x10^4^ cells/well and cultured for 5 days. Spheroids were infected with MG-GFP at MOI 0.1, GFP images were taken 20 hrs post infection and fluorescence measured by the Cytation 5 Imaging Plate Reader (BioTek). Viral replication was measured from supernatants collected 24 hrs post infection and viability determined at 48 hrs by 3D CellTiter-Glo assay following the manufacturer’s instructions (Promega).

### Enzyme-linked immunosorbent assay (ELISA)

The production of human IFNα (Mabtech), IL-28, IL-29, IP-10 (R & D Systems), IL-10 and TNFα (BD Biosciences) and murine IFNγ (R&D Systems) in cell free supernatant was determined using matched-paired antibodies according to the manufacturer’s instructions. Optical density absorbance readings were determined using a Thermo Multiskan EX plate reader, at 405 nm absorbance.

### T2 stability assay

T2 cells were seeded at 1x10^6^/mL with 10ug/mL MG-1 peptide’s (TLLKATPA, RLGPTPPML, SLIQDVERI, GLSKNPVEL, QMTPDVYKV, VLNAPPVQM, KLMDGLINQC, YLYAALKEI, YLLGLYRV; all JPT) cultured for 10 mins at 37°C, then incubated at room temperature overnight. T2 cells were then cultured at 37°C for 2 hrs. T2 cells were then cell surfaced stained for HLA-A2-PB450 or isotype control.

### Cell Surface Phenotyping

Cell surface expression of indicated markers were quantified by flow cytometry. Briefly, cells were harvested, washed in FACS buffer (PBS; 1% (v/v) FCS; 0.1% (w/v) sodium azide), incubated for 30 mins at 4°C with specific antibodies or matching isotype controls. Cells were wash with FACS buffer and then fixed with 1% PFA (1% (w/v) paraformaldhyde in PBS) and stored at 4°C prior to acquisition. Flow cytometry analysis was performed either using BD LSR II flow cytometer Cytek Aurora Spectral cytometer or CytoFLEX S and analysis carried out using either FlowJo software version 0.9 or CytExpert software version 4.3.

### Intra-Cellular Staining

Cells were cell-surfaced stained and fixed overnight with 1% PFA prior to permeabilisation with 0.3% Saponin (Sigma-Aldrich Ltd) for 15mins at 4°C. Cells were wash with 0.1% Saponin, incubated with specific antibodies or matched isotype controls for 30 mins at 4°C. Cells were wash with PBS and flow cytometry analysis was performed immediately using the CytoFLEX S.

### Flow Cytometry Antibodies

*Human: CD11c APC-Vio770 (MJ4-27G12, Miltenyi Biotec), CD14-PerCP (TUK4, Miltenyi Biotec) CD86-PE-Cy7 (2331, BD biosciences), CD80-PE (L307.4, BD biosciences) HLA-ABC-VioBlue (REA230, Miltenyi Biotec), HLA-DR/DP/DQ-FITC (Tu39, BD biosciences), CD3-PerCP (SK7, BD biosciences), CD8-APC (RPA-T8, BD biosciences) IFNg-BV421 (4S.B3, BD biosciences),* HLA-A2-PB450 (BB7.2). Mouse IgG1, κ Isotype Control (PE/ FITC/PerCP/PE-Cy7/APC) (MOPC-21, *BD biosciences*), Mouse IgG2a, κ Isotype Control (FITC/PE) (G155-178, *BD biosciences*), REA Control-VioBlue (REA293, Miltenyi Biotec) and Mouse IgG2bk-PB450 (MPC-11). *Mouse:* CD3 Per-CP (17A2), CD4 BUV395 (GK1.5 BD Biosciences), CD8a BV785 (53-6.7 BD Biosciences), CD8a BV650 (53-6.7) TCRb BUV737(H57-597 BD Biosciences), FOXP3 GFP (), CD25 AF700 (PC61) PD-1 PE-Cy7 (29F.1A12), PD-1-APC (JK3) CXCR5-PeCy7 (DPRCL5) OX40-BV711 (OX-86), all purchased from Biolegend unless stated otherwise.

### Cytotoxic T cell priming assay

Tumour spheroids were infected with MG1 at a MOI 0.1, then cultured with iDC at a 3:1 tumour:iDC ratio for 24 hrs. iDC were collected, washed in PBS and then cultured with autologous PBMC at a 30-40:1 PBMC:iDC ratio for 7 days. Where applicable RNA was collected from week 1 CTL then re-stimulated and cultured for a further 7 days. Primed CTL (week 2) were then harvested, RNA collected, used peptide recall assay or ^51^CR killing assay.

### Peptide Recall assay

To measure peptide specific CTL responses autologous CD14+ cells were incubated with either PMEL, TYR or Mart-1/MLANA PepTivator peptide pools (15-mer peptide sequences with 11 amino acids overlap, Miltenyi Biotec) for 60 min at 37°C, according to the manufacturer’s instructions. Autologous CD14+ cells with or without peptide labelling were then co-cultured with CTL for 60 min at 37°C, Brefeldin A (1:1000, BioLegend) and CD8-APC were then added to cultures and incubated for a further 4 hrs at 37°C. Cells were fixed and then intra-cellular IFNγ-BV421 staining performed prior to analysis by flow cytometry.

### ^51^Cr release assay

Week 2 CTL were cultured with 10ug peptide loaded T2 cells (or unloaded control) at 50:1 E:T ratio for 4 hrs (cells were then pelleted by centrifugation and 50µl of supernatant was transferred to scintillation plates (Perkin-Elmer) prior to analysis using a Wallac Jet 1459 Microbeta scintillation counter and Microbeta Windows software (Perkin-Elmer). Percentage lysis was determined using the following calculation:

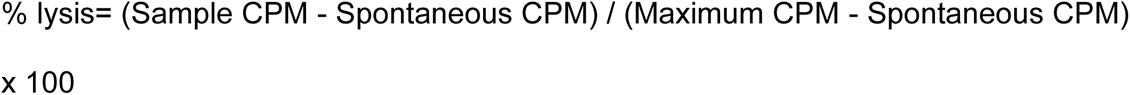

### TCR sequencing

PBMCs from two different donors (D1 and D2) treated under two different conditions (mock and MG1) and two timepoints (week-1 and week-2) were used for TCR sequencing. The sequencing was performed in triplicates. RNA was extracted using TriZol following manufacturer’s instruction (Invitrogen). RNA quantification and quality analysis was performed using Qubit Fluorometry HS kit (Thermo Fisher Scientific) and TapeStation (Thermo Fisher Scientific), respectively, according to manufacturer’s instructions. Bulk RNA-based sequencing of the CDR3β chain was done using a multiplex PCR protocol as previously described^28^. Starting material was 500 ng of RNA diluted in 8 uL of RNase free water. Briefly, this method is multiplex PCR-based and uses 38 primers against the TCR V genes, incorporating unique molecular identifiers (UMIs) to allow the quantification of TCR clones and for the correction of amplifications biases and sequencing errors. Pooled libraries were sequenced according to the Illumina MiniSeq protocol in high output mode (2x150 bp paired end), typically yielding a coverage of 150K reads per sample. The CDR3β extraction and quantification was performed using a computational pipeline for TCR analysis available as a suite of Python scripts available at https://github.com/innate2adaptive/Decombinator.

For the analysis, the TCR repertoires from the triplicates were merged based on donor and condition. Week-2 clonotypes were then classified according to their normalised expansion rate. The expansion rate for each TCR was calculated based on the formula:

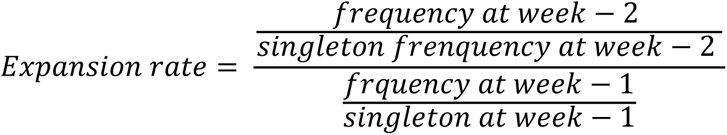

TCRs were classified as expanded if normalised expansion rate was > 1 or contracted if normalised expansion rate was < 1. Each repertoire group was subsequently clustered based on CDR3β amino acid triplet similarity using a kernel matrix and a similarity threshold of 0.65. Normalised cluster count (to the top 500 TCRs) and cluster size were quantified.

### Animals

Five- to six-week-old female C57Bl/6 mice (Charles River, UK) or Nr4a3 Tocky mice (C57Bl/6 background) were used for tumour implantation. These were conducted at the ICR Biological Services Unit and were approved by the ICR local Ethical Review Committee and standards of care were based upon the UKCCCR Guidelines for the welfare and use of animals in cancer research. 4434 murine melanoma tumours were established by 100 µL subcutaneous injection of 4x10^6^ cells respectively into the right flank of each mouse. Tumour measurements were taken twice weekly in three dimensions using Venier callipers and the tumour volume estimated using the formula: length x width x height (mm) x 0.5236. For flow cytometry analysis, tumours were harvested at indicated volumes and were mechanically digested with scissors, then digested in 1ml of a digestion mix containing 25ug/ml Liberase, 250uL/ml DNASEI, 40uL/ml Trypsin at 37C for 10 mins, then RT for 20 mins. Digested contents passed through 70uM cell strainer and washed with FACS buffer.

### RNA sequencing of 4434 tumours

4434 tumours were explanted from animals 48 hrs following treatment and stored in RNALater (AM7020, Thermo Fisher) at -20°C prior to RNA extraction. Samples were homogenised as described previous and RNA extraction performed using RNEasy kit (74104, Qiagen, Germantown, USA) as per manufacturer protocol.

### Bioinformatics analysis

Trimmomatic (v0.39) was used for raw reads trimming, followed by Hisat2 (v2.1.0) alignment software for trimmed reads mapping to Ensemble GRCm38.102 mouse reference genome. Stringtie (v2.1.4) in combination with custom python script were used to generate gene count matrix for all samples. Genes that had less than 1 read per sample were removed from further analysis. Differential expression analysis was performed in R using the Bioconductor package DESeq2 and mMCP counter was applied to estimate immune and stromal cell type signatures for each sample. Over- representation analysis of differentially expressed genes was carried out in using enrichR web-application.

### Virus treatments

A single intra-tumoural dose of PBS or MG-FLUC (1x10^7^PFU) was given when tumours reached the desired volume. For combination experiments 200 ug of isotype control (InVivoMab mouse IgG2b isotype control, clone MPC-11; 2BScientific), or anti-PD-1 (InVivoMab rat anti-mouse PD-1 (CD279), clone- RMP1-14 monoclonal antibody, IgG2a κ; 2BScientific) antibody was administered intra-peritoneally twice a week for a maximum of four weeks or until mice reached humane end point. Tumour measurements were recorded twice weekly.

### Histology and Immunohistochemistry

Tumours from treated mice were dissected and fixed overnight in formalin. These were then processed, embedded and 2 µm sections were prepared on APEX glass slides. Sections from formalin-fixed paraffin-wax embedded tumours were stained with primary rat anti-mouse monoclonal anti-CD8 antibody (eBioscience, 4Sm15as) and goat anti-mouse polyclonal anti-PD-L1 antibody (R&D systems, AF1019). Slides were scanned and imaged using Hamamatsu Nanozoomer (Hamamatsu Photonics).

### Splenocyte recall assay

Spleens were immediately excised from euthanized mice and dissociated *in vitro* to achieve single-cell suspensions. Red blood cells were lysed with ACK lysis buffer for 1 minute. Cells were resuspended at 1 × 10^6^ in glutamine containing RPMI, supplemented with 10% FCS and 1% Pen-Strep. Splenocytes were cultured either alone or with the indicated tumour cells at a ratio of 5:1 (E:T) or Rhabdovirus N peptide (RGYVYQGL). Cell-free supernatants were collected 48 hours later and tested by IFNγ ELISA.

### Statistical Significance

Statistical analysis was carried out with the GraphPad Prism software. Statistical differences among groups were determined using student’s t-test, one-way ANOVA or two-way ANOVA analysis. For survival experiments, the Kaplan-Meier survival curves were compared using log-rank (Mantel-Cox) test. Statistical significance was determined as follows: *\*p<0.05, **p<0.0021, ***p<0.0002* and *\*\*\*\*p<0.0001*.

## List of abbreviations

APC: Antigen presenting cell
HSV: Herpes simplex virus
ICI: Immune checkpoint inhibitors
iDC: Immature dendritic cell
MART-1/MELAN-A: Melanoma antigen recognized by T-cells 1
NCI: National Cancer Institute
Ovs: Oncolytic Viruses
PMEL: Premelanosome
TAA: Tumour associated antigen
TIL: Tumour infiltrating lymphocyte
TIME: Tumour immune microenvironment
TME: Tumour microenvironment
T-VEC: Talimogene laherparepvec
TYR: Tyrosinase

